# A multitask transfer learning framework for novel virus-human protein interactions

**DOI:** 10.1101/2021.03.25.437037

**Authors:** Ngan Thi Dong, Megha Khosla

## Abstract

Understanding the interaction patterns between a particular virus and human proteins plays a crucial role in unveiling the underlying mechanism of viral infection. This could further help in developing treatments of viral diseases. The main issues in tackling it as a machine learning problem is the scarcity of training data as well input information of the viral proteins. We overcome these limitations by exploiting powerful statistical protein representations derived from a corpus of around 24 Million protein sequences in a multi task framework. Our experiments on 7 varied benchmark datasets support the superiority of our approach.

## 1 Introduction

Viral infections most have been increasingly burdening the healthcare systems. Biologically the viral infection involves many protein-protein interactions (PPIs) between the virus and its host. These interactions range from the initial biding of viral coat proteins to the host membrane receptor to the hijacking of the host transcription machinery by viral proteins. In this work we develop a deep learning based computational model for predicting interactions between a novel virus (a completely new one) and human proteins.

One of the key challenges in tackling the current learning task with novel unseen viruses is the *limited training data*. Often, some known interactions of related viruses are used to train supervised models. These data is usually collected by wet lab experiments and are usually too little to ensure generalizability of trained models. In effect, the trained models might overfit the training data and would give inaccurate predictions for the novel virus.

Moreover, viral proteins are substantially different from human or bacterial proteins. They are structurally dynamic so that they cannot be easily detected by common sequence-structure comparison (Requião et al., 2020). Virus protein sequences of different species share only little in common (Eid et al., 2016). Therefore, models trained for other human PPI (Li & Ilie, 2020; Sun et al., 2017; Li, 2020; Chen et al., 2019; Sarkar & Saha, 2019) or for other pathogen-human PPI (Sudhakar et al., 2020; Mei & Zhang, 2020; Dick et al., 2020; Li et al., 2014; Guven-Maiorov et al., 2019; Basit et al., 2018)(for which more data might be available) cannot be directly used for predictions for novel viral-human protein interactions.

While for human proteins, features related to their function, semantic annotation, domain, structure, pathway, etc. can be extracted from public databases, such information is not readily available for viral proteins. The only reliable source of viral protein information is its amino acid sequence. *Learning effective representations* of the viral proteins is thus an important step towards building the prediction model. Heuristics such as K-mer composition usually used for protein representations are bound to fail as it is known that viral proteins with completely different sequences might show similar interaction patterns.

Other existing works also employed additional features to represent viral proteins such as protein functional information (or GO annotation) (Wang, 2020), proteins domain-domain associations information as in (Barman et al., 2014), protein structure information as in (Lasso et al., 2019; Guven-Maiorov et al., 2019), and the disease phenotype of clinical symptoms as in (Wang, 2020). A major limitation of these approaches is that they cannot generalize to novel viruses where such information is not available or lack experimentally supported evidence.

In this work we tackle the above limitations by exploiting powerful statistical protein representations derived from a corpus of around 24 Million protein sequences in a multitask framework. Noting the fact that virus tends to mimic humans towards building interactions with its proteins, we use the prediction of human PPI as a side task to further regularize our model and improve generalization. Our large scale experiments on a number of datasets showcase the superiority of our approach.

## 2 Our approach

The schematic diagram of our proposed model is presented in Figure 1. We use human and virus raw protein sequences as input. As side or domain information we use human protein-protein interaction network of around 20,000 proteins and over 22M interactions from STRING (Szklarczyk et al., 2015) database.

**Figure 1:**
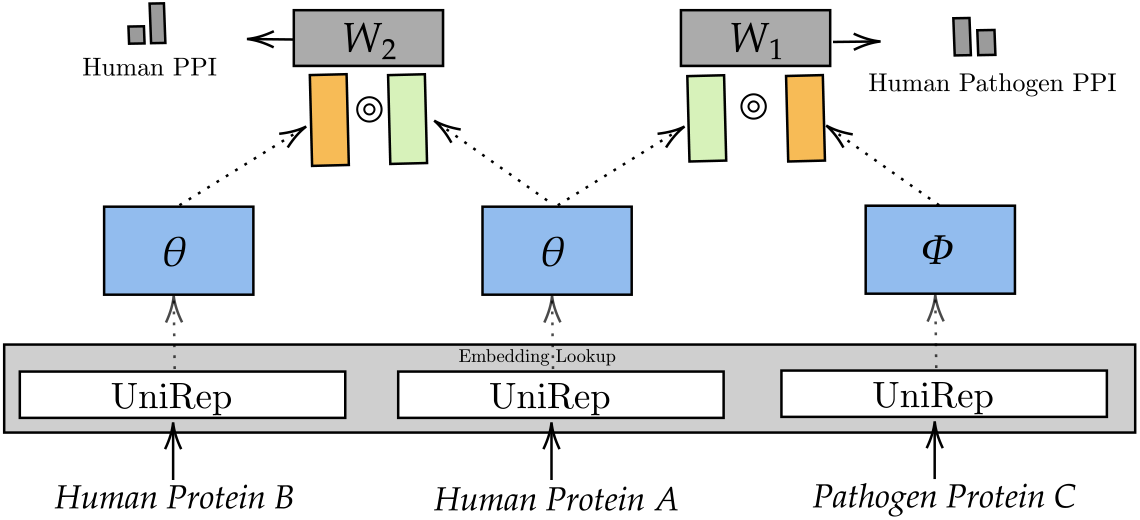
Multitask Transfer (MTT) model for pathogen-human PPI.

We note that the protein sequence determines the protein’s structural conformation (fold), which further determines its function and its interaction pattern with other proteins. However, the underlying mechanism of the sequence-to-structure matching process is very complex and cannot be easily specified by hand crafted rules. Therefore, rather than using handcrafted features extracted from amino acid sequences we employ the pre-trained UniRep model (Alley et al., 2019) to generate latent representations or protein embeddings. The protein representations extracted from UniRep model are empirically shown to preserve fundamental properties of the proteins and are hypothesized to be statistically more powerful and generalizable than hand crafted sequence features.

We further fine-tune these representations by training 2 simple neural networks (single layer MLP with ReLu activation) using an additional objective of predicting human PPI in addition to the main task. We use Logistic Regression networks to predict likelihood of having interaction between virus-human proteins or human-human proteins. The two networks’ parameters are not shared among tasks allowing them to extract more task-specific representation.

The rationale behind using human PPI task is that viruses have been shown to mimic and compete with human proteins in their binding and interaction patterns with other human proteins (Mei & Zhang, 2020). Therefore, we believe that the patterns learned from the human interactome (or human PPI network) should be a rich source of knowledge to guide our virus-human PPI task and further helps to regularize our model.

Let Θ, Φ denote the set of learnable parameters corresponding to representation tuning components, i.e., the Multilayer Perceptrons (MLP) corresponding to the virus and human proteins, respectively. Let **W**_1_, **W**_2_ denote the two learnable weight matrices (parameters) for the logistic regression modules for the virus-human and human-human PPI prediction tasks. We use *V H*, and *HH* to denote the training set of virus-human, human-human PPI, correspondingly.

We use binary cross entropy loss for virus-human PPI predictions as given below

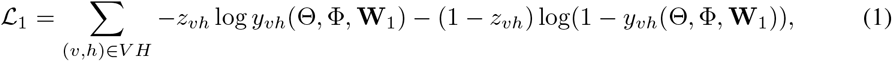

where variables *z*_*vh*_ is the corresponding binary target variable and *y*_*vh*_ is the predicted probability of virus-human PPI or the output of the Logistic regression (LR) module. The input to the LR module is the element wise product of fine-tuned representations (output of the MLP) of virus and human protein.

For human PPI, the target variables (*z*_*hh ′*_) are the normalized confidence scores which can be interpreted as the probability of observing an interaction. We use binary cross entropy loss as below where *y*_*hh′*_ is the element wise product of fine-tuned representations (output of the second MLP) of human and human protein.

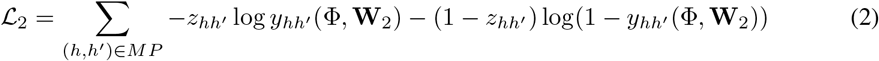

We use a linear combination of the two loss functions to train our model, i.e., ℒ= ℒ_1_ + α ·*ℒ*_2_,

where α is the human PPI weight factor. We set it to 10^*−*3^ in our experiments.

## 3 Experimental Evaluation

We compare our method with following six baseline methods and two simper variants of our model.

1. Generalized (Zhou et al., 2018): It is a generalized SVM model trained on hand crafted features extracted from protein sequence for the novel virus-human PPI task.
2. Hybrid (Deng et al., 2020): It is a complex deep model with convolutional and LSTM layers for extracting latent representation of virus and human proteins from their input sequence features and is trained using L1 regularized Logistic regression.
3. DOC2VEC (Yang et al., 2020): It employs the doc2vec (Le & Mikolov, 2014) approach to generate protein embeddings from the corpus of protein sequences. A random forest model is then trained for the PPI prediction.
4. MotifTransformer(Lanchantin et al., 2020): It first generates protein embeddings using supervised protein structure and function prediction tasks. Those embeddings were later passed as input to a an order-independent classifier to do the PPI prediction task.
5. DeNovo(Eid et al., 2016): It trained a SVM classifier on a hand crafted feature set extracted from the K-mer amino acid composition information using a novel negative sampling strategy.
6. Barman(Barman et al., 2014): It used a SVM model trained on feature set consisting of the protein domain-domain association and methionine, serine, and valine amino acid composition of viral proteins.
7. 2 simpler variants of MTT: Towards ablation study we evaluate two simpler variants: (i) SingleTask Transfer (STT), which is trained on a single objective of predicting pathogen-human PPI and (ii) NaiveBaseline, which is a Logistic regression model using concatenated human and pathogen protein UniRep representations as input.

### 3.1 Benchmark Datasets and Results

We evaluate our approach on 7 benchmark datasets. As several of our competitors do not release their code, we use the reported performance scores (using the same evaluation metrics) in the original papers giving them full advantage. Besides, as many of the methods use hand crafted features which might not be available for other benchmark datasets not evaluated in their original papers. Detailed data statistics can be found in the Appendix A.1.

#### Novel Viral-Human PPI

We use the benchmark datasets for human H1N1 and human Ebola viruses as released by Zhou et al. (2018). The dataset is prepared for testing predictions for a novel virus. The known PPIs between virus and human were retrieved from four databases: APID, IntAct, Metha, and UniProt. The training data for the human-H1N1 dataset includes PPIs between human and all viruses except H1N1. Similarly, the training data for the human-Ebola dataset includes PPIs between human and all viruses except Ebola. The statistics for both datasets are presented in Table 4 in the Appendix. The results (Area under curve (AUC) and Area under Precision Recall curve (AUPR) scores) are given in Table 1.

**Table 1:** Comparison on novel virus-human PPI prediction task. “-” denotes that the corresponding score is not reported in the original paper.

| MODEL | H1N1 |  | EBOLA |  |
| --- | --- | --- | --- | --- |
|  | AUC | AUPR | AUC | AUPR |
| GENERALIZED | 0.886 | - | 0.867 | - |
| HYBRID | 0.937 | - | - | - |
| MOTIFTRANSFORMER | 0.945 | 0.948 | 0.968 | 0.974 |
| DOC2VEC | 0.817 | 0.542 | - | - |
| MULTITASK TRANSFER(MTT) (our) | **0.957** | **0.966** | **0.976** | **0.981** |
| SINGLETASK TRANSFER(STT) (our) | 0.950 | 0.962 | 0.963 | 0.974 |
| NAIVE BASELINE (our) | 0.834 | 0.806 | 0.893 | 0.870 |

**Table 2:** Comparison on datasets with rich feature information. SN, SP, ACC refer to *Sensitivity, Specificity*, and *Accuracy*, respectively. “-” denotes that the corresponding score is not reported in the original paper.

| MODEL | DENOVO SLiM |  |  |  | BARMAN’S DATASET |  |  |  |
| --- | --- | --- | --- | --- | --- | --- | --- | --- |
|  | SN | SP | ACC | AUC | SN | SP | ACC | AUC |
| GENERALIZED | 80.00 | 88.94 | 84.47 | 0.897 | 76.14 | 83.77 | 79.95 | 0.858 |
| DENOVO Eid et al. (2016) | 82.59 | 81.65 | 83.53 | - | - | - | - | - |
| MTT (our) | **88.00** | **87.76** | **87.88** | **0.955** | 90.05 | 89.57 | 89.81 | **0.958** |
| STT (our) | 86.12 | 85.88 | 86.00 | 0.941 | **90.14** | **89.66** | **89.9** | 0.957 |
| NAIVE BASELINE (our) | 84.00 | 83.76 | 83.88 | 0.885 | 74.20 | 73.72 | 73.96 | 0.809 |

#### Viral-Human PPI prediction on Datasets with Rich Viral information

We use the datasets from DeNovo(Eid et al., 2016) and Barman (Barman et al., 2014) studies. DeNovo’s SLiM dataset encapsulated viral proteins based on presence of Short Linear Motif (SLiM) (short recurring protein sequences with specific biological function). Barman’s dataset was retrieved from Virus-MINT database by removing interacting protein pairs that did not have any “InterPro” domain hit. Barman dataset is evaluated using 5-fold cross validation using the original data splits.

#### Additional results on novel bacteria-human PPI prediction

We further demonstrate our model effectiveness on the novel bacteria-human PPI prediction task. We compare our method with Denovo on the three datasets for three human bacteria: Bacillus anthracis (B1), Yersinia pestis (B2), and Francisella tularensis (B3), obtained from (Eid et al., 2016). The results are shown in Table 3. MTT clearly outperforms the baseline method (Denovo).

**Table 3:** Comparison for the novel bacteria-human PPI prediction task. SN, SP, ACC refer to *Sensitivity, Specificity*, and *Accuracy*, respectively.

| MODEL | BACILLUS ANTHRACIS |  |  | YERSINIA PESTIS |  |  | FRANCISELLA TULARENSIS |  |  |
| --- | --- | --- | --- | --- | --- | --- | --- | --- | --- |
|  | SN | SP | ACC | SN | SP | ACC | SN | SP | ACC |
| DENOVO | **94** | 97.2 | 96.42 | 94.8 | 98.3 | 97.47 | 94.9 | 98.3 | 97.32 |
| MTT(our) | 93.46 | **97.83** | **96.74** | **96.93** | **98.99** | **98.49** | **98.22** | **99.27** | **98.98** |

**Table 4:** Novel virus-human PPI benchmark datasets’ statistics. |*E*^+^| and | *E*^*−*^| refer to the numebr of positive and negative interactions, respectively. *V* ^*h*^ and *V* ^*v*^ are the number of human proteins and viral proteins.

|  | TRAINING DATA |  |  |  | TESTING DATA |  |  |  |
| --- | --- | --- | --- | --- | --- | --- | --- | --- |
| | $\vert E^+\vert $ | $\vert E^-\vert $ | $\vert V^h\vert $ | $\vert V^v\vert $ | $\vert E^+\vert $ | $\vert E^-\vert $ | $\vert V^h\vert $ | $\vert V^v\vert $ |
| HUMAN-H1N1 | 10,858 | 10,858 | 7,636 | 641 | 381 | 381 | 622 | 11 |
| HUMAN-EBOLA | 11,341 | 11,341 | 7,816 | 659 | 150 | 150 | 290 | 3 |

### 3.2 Discussion and Future Work

Our methods shows superior performance on a wide range of tested datasets. Note that this is despite the fact that each of our baselines have been proposed to exploit certain specific kind of information which was in the first place used to construct the dataset. MTT also outperforms its simpler variants developed with single task objective. Note that our naive baseline which directly trains a logistic regression classifier with pretrained embeddings already outperforms several methods. This points to the superiority of these representations as compared to hand-crafted features. As future work We will enhance our multi task approach by incorporating more domain information as well as exploiting more sophisticated multi task model architectures.

## A Appendix

### A.1 Data Description

In this section we provide further details and statistics of the 7 benchmark datasets.

#### A.1.2 Data for novel-human PPI

We use the benchmark datasets for human H1N1 and human Ebola viruses as released by (Zhou et al., 2018). The dataset is prepared as follows and is suitable for testing predictions for novel viruses.

The known PPIs between virus and human were retrieved from four databases: APID, IntAct, Metha, and UniProt. In total, there are 11,491 known PPIs between 246 viruses and human. The training data for the human-H1N1 dataset includes PPIs between human and all viruses except H1N1. Similarly, the training data for the human-Ebola dataset includes PPIs between human and all viruses except Ebola. The testing data for the human-H1N1 dataset contains PPIs between human and 11 H1N1 viral proteins. Likewise, the testing data for the human-Ebola dataset contains PPIs between human and 3 Ebola proteins.

The statistics for those datasets are presented in Table 4.

#### A.1.2 Data with Rich Viral information

Denovo SLiM dataset (Eid et al., 2016)Short Linear Motif (SLiM) are short, recurring patterns of protein sequences that have a biological function. They are believed to mediate protein-protein interaction (Diella et al., 2008; Neduva & Russell, 2006). Therefore, sequence motifs can be a rich feature set for virus-human PPI prediction tasks.

Denovo’s SLiM testing set contained 425 positives and 425 negative PPIs. Since SLiM information is only available for a very limited number of viral proteins, this dataset gave more advantages for heuristics-based methods. The training data consisted of the remaining PPI records in VirusMentha database (Calderone et al., 2015) (except the ones in the testing data). In the end, the training data comprised of 1590 positive and 1515 negative records with known SLiM information and 3430 positives and 3219 negatives without SliMs information.

Barman’s dataset (Barman et al., 2014). The dataset was retrieved from VirusMINT database (Chatr-Aryamontri et al., 2009). Interacting protein pairs that did not have any “InterPro” domain hit were removed. In the end, the dataset contained 1035 positive and 1,035 negative interactions between 160 virus proteins of 65 types and 667 human proteins. 5-Fold cross-validation was then employed to test each method’s performance.

#### A.1.3 Data for novel Bacteria-Human PPI (Eid et al., 2016)

The data was first collected from HPIDB (Ammari et al., 2016). B1 belongs to a bacterial phylum different from that of B2 and B3, while B2 and B3 share the same class but differ in their taxonomic order. B1 has 3057 PPIs, B2 has 4020, and B3 has 1346 known PPIs. A sequence-dissimilarity-based negative sampling method was employed to generate negative samples. For each bacteria protein, ten negative samples were generated randomly. Each of the bacteria was then set aside for testing, while the interactions from the other two bacteria were used for training. In the end, we have three datasets. For simplicity, we use the name of the bacteria in the testing set as the name of the dataset. The statistics for those three datasets are presented in table 5.

**Table 5:** Our novel bacteria-human PPI benchmark datasets’ statistics. |*E*^+^| and |*E*^*−*^ |refer to the number of positive and negative interactions, respectively. |*V* ^*h*^ |and |*V* ^*b*^ |are the number of human proteins and bacteria proteins.

|  | TRAINING DATA |  |  |  | TESTING DATA |  |  |  |
| --- | --- | --- | --- | --- | --- | --- | --- | --- |
| | $\vert E^+\vert $ | $\vert E^-\vert $ | $\vert V^h\vert $ | $\vert V^b\vert $ | $\vert E^+\vert $ | $\vert E^-\vert $ | $\vert V^h\vert $ | $\vert V^b\vert $ |
| BACILLUS ANTHRACIS | 5366 | 15590 | 1559 | 2674 | 3057 | 9440 | 944 | 1705 |
| YERSINIA PESTIS | 4403 | 12880 | 1288 | 2278 | 4020 | 12150 | 1215 | 2147 |
| FRANCISELLA TULARENSIS | 7077 | 21590 | 2159 | 3041 | 1346 | 3440 | 344 | 1023 |

